# Large effects and the infinitesimal model

**DOI:** 10.1101/2023.07.20.549972

**Authors:** Todd L. Parsons, Peter L. Ralph

## Abstract

The infinitesimal model of quantitative genetics relies on the Central Limit Theorem to stipulate that under additive models of quantitative traits determined by many loci having similar effect size, the difference between an offspring’s genetic trait component and the average of their two parents’ genetic trait components is Normally distributed and independent of the parents’ values. Here, we investigate how the assumption of similar effect sizes affects the model: in particular, if the tail of the effect size distribution is polynomial with exponent *α <* 2, then sums of effects should be well-approximated by a “stable distribution”, and find tail exponents between 1 and 2 in effect sizes estimated by genome-wide association studies of many human disease-related traits. We show that the independence of offspring trait deviations from parental averages in many cases implies a Gaussian distribution, suggesting that non-Gaussian models of trait evolution must explicitly track the underlying genetics, at least for loci of large effect. We also characterize possible limiting trait distributions of the infinitesimal model with infinitely divisible noise distributions, and compare our results to simulations.

## 1 Introduction

Much of the work to understand the evolution of quantitative traits in sexual species has focused on models in quantitative genetics that describe evolution of trait values without explicitly representing underlying genetic variation, but in a manner deriving from our understanding of how genetic variation works in practice. Such models are therefore interestingly intermediate between mechanistic models of genetics and purely phenomenological. One concrete model of heritable trait evolution is the *infinitesimal model*, which assumes a simple, explicit form for how the genetic component of offsprings’ traits are determined by their parents: they are Normally distributed, with the mean for each offspring equal to the average of the parents’ values, and a covariance independent of the parental trait values (but that depends on how they are related). Barton et al. [2017] give a history of the model and a formal proof of conditions under which this would be a good approximation to reality, which they extended to include dominance in Barton et al. [2022].

As suggested by its name, under the infinitesimal model the effects of any given genetic variant on the trait is small (indeed, vanishing), and selection-driven trait shifts are mediated by small changes in frequencies of many alleles of small effect. However, empirical understanding of the genetic basis of trait variation in real-world species often finds a significant portion of heritable variation explained by variants at only a few loci. There are many possible explanations for this observation, including selection or publication bias, filters for larger-effect variants due to spatially varying selection [Barton, 1987, Yeaman and Otto, 2011, Yeaman and Whitlock, 2011], and/or grouping together of concordant alleles by linkage or inversions [Kirkpatrick and Barton, 2006]. However, it seems clear that in at least those cases where we understand the underlying molecular mechanisms, mutations of large effect can be important in practice. The consensus in the field seems to be that traits exist on a spectrum between monogenic and polygenic, and that many - if not most - lie in the intermediate region, where heritable trait variation is due both to segregating large- and small-effect variants. Indeed, some sensible verbal models of the biological mechanisms by which a genetic change comes to have an effect on an organism’s trait describe a hierarchy or range of “proximities” to the trait, by which changes to more “nearby” genes have the possibility of causing larger changes in the trait [Kopp and McIntyre, 2012, Boyle et al., 2017]. This suggests a mixture model for the effect size distribution: perhaps the distribution of effect sizes within each gene is Normal, but with a standard deviation that varies with “proximity” to the trait; such mixtures can easily have large proportions of extreme values.

The Gaussian distribution appears thanks to “the” Central Limit Theorem, i.e., the universality of adding up a great many very small and independent things. However, there is a wider class of Central Limit Theorems, which describe the distributions of sums of a great many independent and exchangeable things: the *stable laws* [Gnedenko and Kolmogorov, 1968]. Roughly speaking, these say that if we add together *n* independent copies of a random variable *X* for which ℙ {*X > t*} is proportional to *t*^−*α*^ for some 0 *< α <* 2, then – almost regardless of the details – the result, scaled by *n*^1*/α*^, is well-approximated by a universal form, an *α*-stable distribution (and the approximation becomes exact, as *n* → ∞). In such ensembles, single large entries can be important: concretely, the largest of the *n* copies will be of order *n*^1*/α*^, so that e.g., for *α* = 1 a positive proportion of the sum of *n* entries will be contributed by a single one.

This suggests exploring whether stable distributions might reasonably stand in for the Gaussian in the infinitesimal model. In this paper, we make preliminary explorations in that direction: Do the results of Barton et al. [2017] carry over when the distribution of effect sizes falls in the domain of attraction of a stable law? The immediate answer to this will be “no” – in fact, we will see that independence of offsprings’ deviations and midparent values *implies* a Gaussian distribution in some sense. This independence is a key reason why the infinitesimal model is tractable, and so a conclusion is that additive models with non-Gaussian effects may not be well-approximated by “trait-only” models that do not explicitly represent underlying genetic variation. Having thus shown that “infinitesimal models with non-Gaussian noise” is on weak footing as a model of reality, we then go on to analyze exactly these models. (As motivation, we have not shown that such models are *not* good approximations to reality, and it is interesting and perhaps informative to understand how the noise distribution affects results.)

Stable distributions are by no means new to population biology. For instance, it is a long-standing question how often the motions of organisms are better modeled by essentially Brownian random walks or by Lévy processes [Benhamou, 2007] and a growing literature studies the effects of such power law distributions on spatial genetic patterns and evolutionary dynamics [e.g., Paulose et al., 2019, Smith and Weissman, 2023]. More directly analogous to the problem we study here, Landis et al. [2012] found that an *α*-stable process (with *α* ≈ 3*/*2) fit the evolution of log endocranial volume on the primate phylogeny better than a Brownian model. It is also important to note that the infinitesimal model implies a Gaussian distribution for the difference between an offspring’s trait and their parents’ average, *not* for the entire population. The question of when the trait distribution in an evolving population might be modeled as a Gaussian has been discussed extensively [e.g., Turelli and Barton, 1994, Débarre et al., 2015].

The ways in which genetic variants affect an organisms’ trait can be complex, and interactions between alleles, between loci, and with the environment are often important in practice. In fact, many traits claimed to be genetically determined in fact only appear heritable through non-genetic means, e.g., systematic inequalities in resource distribution partitioned by class or race. In this paper, we follow the literature in working with the relatively simple purely additive case of a trait that can be decomposed into a sum across loci of the contributions of the alleles from each locus, even omitting an independent effect of “the environment”. However, we emphasize that such models should not always be the default for understanding biological variation, especially for human traits.

### Notation

We generally follow the notation of Barton et al. [2017], although in a simpler situation, as we do not consider full pedigrees but rather only the haploid offspring of a single set of haploid parents (as produced, for instance, by a round of meiosis in a diploid stage), and ignore non-genetic contributions to the trait; so, below we say “trait” to mean “genetic contribution to the trait”.

Suppose that the focal individual’s trait is

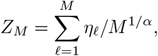

where *η*_*ℓ*_ is the contribution of the allele carried at the *ℓ*^th^ locus. The effects of the alleles carried by the two parents are 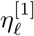 and 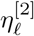, respectively, and their traits are 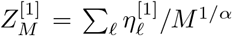 and 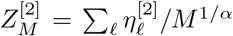. We assume the loci are unlinked and inheritance is fair, so letting *X*_*ℓ*_ be a family of Bernoulli(1/2) random variables that are independent of everything else,

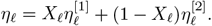

What we will call the *Mendelian sampling term* (or occasionally *Mendelian noise*), *R*_*M*_, is the offspring’s trait minus the midparent value, 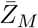:

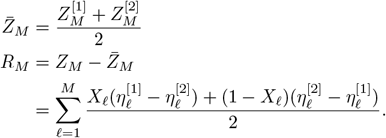

Note in particular that the contribution of locus *ℓ* to *R*_*M*_ is 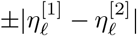.

## 2 Empirical observations

It is mathematically intriguing to speculate what effect a power law effect size distribution for would have on the induced evolutionary dynamics. But, is there any evidence that non-Gaussian effects are important in practice? In this paper we focus primarily on the theory, but first take a brief and imperfect look at what some data have to say on the topic. To do this, we downloaded the GWAS results provided by Abbott et al. for 7,221 phenotypes from 361,194 human individuals from the March 2018 release of the UK Biobank. After filtering (see details below) we were left with 695 binary illness-related phenotypes. Taken at face value, these provide for each phenotype a set of estimates of the additive effect of the alternative allele at a set of single nucleotide polymorphisms (SNPs) on the phenotype relative to the reference allele. The phenotypes we consider were all binary and coded as 0/1, so estimates could be thought of as additive adjustments to the risk (so, effects are *not* on a logit scale). These estimates were obtained as the coefficient for the SNP in a simple linear model fit to each phenotype for each SNP, with the first 20 principal components and all combinations of age, age squared, and inferred sex as covariates; we considered effects of those SNPs with a reported *p*-value less than 10^−8^. There are a great many possible issues with these data; however, the results seem consistent across many SNPs and phenotypes, and we know of no reason that possible methodological artifacts (e.g., confounding by uncorrected variables) would specifically affect the decay of the tail of effect sizes.

Since the mendelian sampling term is a sum of terms of the form

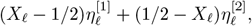

where 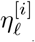 is the effect of the allele in parent *i* at locus *ℓ*, the appropriate central limit theorem for the sum is determined by the tail behavior of |(*X* − 1*/*2)*η* |= |*η*| */*2, where *X* is Bernoulli(1/2) and *η* is the effect of a uniformly chosen allele at a uniformly chosen locus. So, we’d like to know if ℙ {|*X*| *>* 2*t*} is proportional to *t*^−*α*^, i.e., the value of

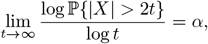

if the limit exists. Let *p*_*ℓ*_ be the minor allele frequency at the *ℓ*^th^ locus; since the data we have assigns effect 0 to the major allele, *η*_*ℓ*_ = 0 with probability 1 − *p*_*ℓ*_. Therefore, if we have estimated effects for *M* SNPs with minor allele frequency *p*_*ℓ*_ and effect size *e*_*ℓ*_ for the minor allele, for 1 ≤ *ℓ* ≤ *M*, then

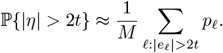

So, to obtain a per-trait estimate of *α*, we plot 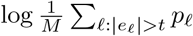 against log *t* for a sequence of values of *t*, and fit a linear model to the tail (which we take to be all values of *t* for which the number of SNPs with absolute effect above *t* is between 20 and 10% of all SNPs). Examples are shown in Figure 1.

**Figure 1.**
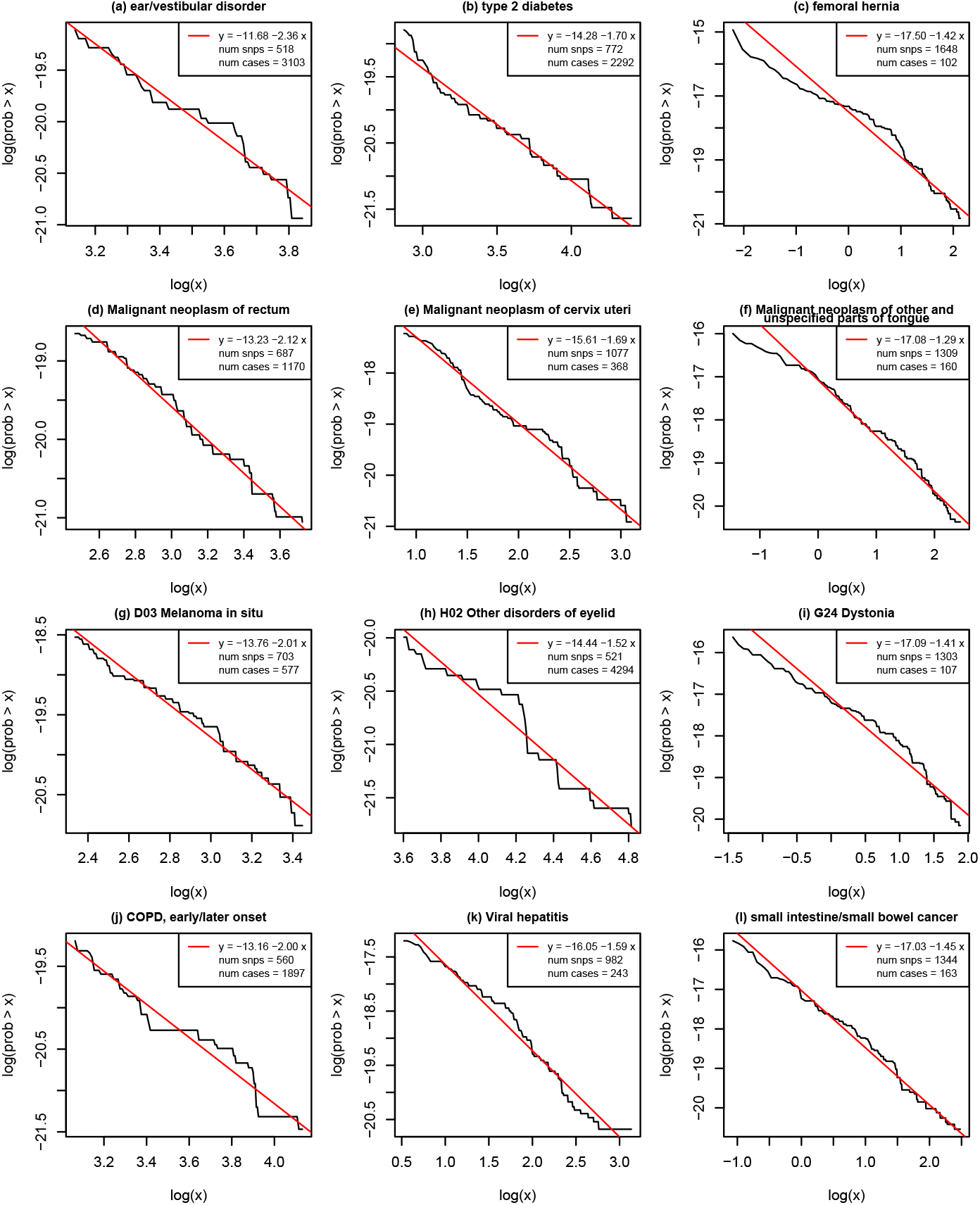
Tail distributions of frequency-weighted effects for twelve randomly chosen phenotypes. In each, for a phenotype with *N* SNPs, effect sizes *e* and frequencies *p*, the black line shows 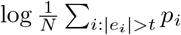 (vertical axis) against log *t* (horizontal axis); the red line shows the least-squares linear fit; each plot only covers the 10% largest-effect SNPs. Legends show th5e coefficients of the red line; the slope on ‘x’ is *α*, the estimated exponent; “num snps” is the number of SNPs for the phenotype with *p*-value less than 10^−8^, and “num cases” is the reported number of the roughly 360,000 individuals recorded as having the listed phenotype.

We applied this procedure to results from models fit on both sexes, after removing phenotypes related to non-medical traits (e.g., employment, drug treatments, and dietary choices). This left us with 1,108 phenotypes for which we had enough data (e.g., enough significant SNPs) to estimate the tail exponent. We then restricted to phenotypes having non-missing numbers of “cases” and “controls” and a sample size of at least 300,000 (i.e., not more than 61,194 of 361,194 missing phenotypes), and removed the 65 remaining phenotypes that are common (more than 10,000 cases), as these often showed very different patterns (e.g., effect size distributions). This left us with 695 phenotypes; plots including the 413 phenotypes removed by these last steps are provided in Supplementary Figure S1.

Resulting values of the tail exponent *α* are shown in Figure 2. Estimated tail exponents range from about 1 to 2.5, and 28.9% had estimated values less than 1.5. A concern is that “noisier” phenotypes, for which estimated effect sizes are less reliable, might lead to larger estimated tail exponents; for this reason, we looked for an association between tail exponent and two proxies for power, number of SNPs (i.e., number of SNPs with *p*-value less than 10^−8^) and number of cases. Interestingly, the relationship is nonmonotonic, with exponents closer to *α* = 2 for phenotypes with around 1000 cases, and lower exponents for both rarer and more common phenotypes. (A similar pattern is seen for “number of SNPs”, but this may be due to association with power and hence number of cases.) This may indicate a statistical artifact, or it may be a result of biological differences in genetic architecture between more and less common phenotypes.

**Figure 2.**
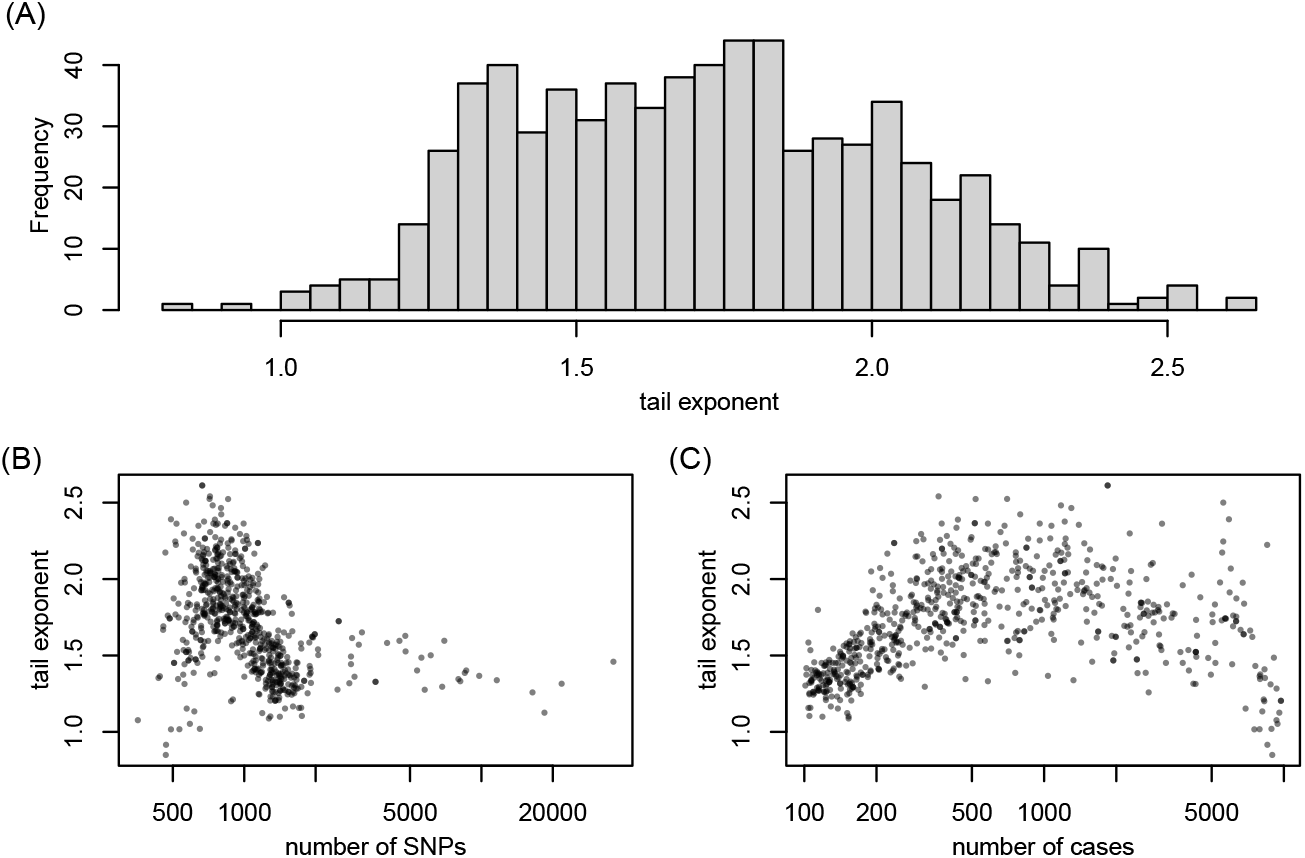
Estimated values of the tail exponent, *α*, for 775 illness-related binary phenotypes: **(A)** distribution of values; and plotted against **(B)** number of SNPs, and **(C)** number of cases.

## 3 Independence and the Gaussian assumption

We know from Barton et al. [2017] that if the 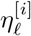 are independent with mean zero and 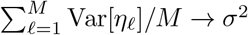, then with *α* = 2, the traits (*Z*_*M*_, 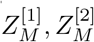) converge in distribution to a multivariate Gaussian distribution (*Z, Z*^[1]^, *Z*^[2]^) for which *Z* − (*Z*^[1]^ + *Z*^[2]^)*/*2 is independent of *Z*^[1]^ and *Z*^[2]^.

Perhaps surprisingly, there is a converse – in at least some situations, independence of the Mendelian sampling term implies a Gaussian distribution. To see this, divide the parental loci into groups according to the results of inheritance, defining

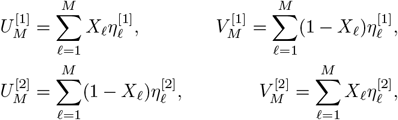

so that 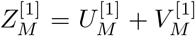 and 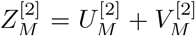, while 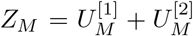 is the transmitted portion of the parental genomes, and we can define 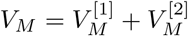 to be the untransmitted portion. Furthermore,

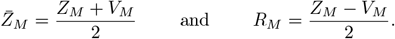

Now, note that if inheritance (the *X*_*ℓ*_) is independent of the per-locus effects 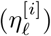, the latter are all independent of each other, and all effects at a given locus are drawn from the same distribution, then the inherited allele, 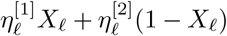 is independent of (and has the same distribution as) the non-inherited allele, 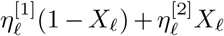 they are simply two independent draws from the distribution of effects at locus *ℓ*. Therefore, the inherited alleles that sum to *Z*_*M*_ are independent of the non-inherited alleles that sum to *V*_*M*_.

The Kac-Bernstein theorem states that if two independent random variables have the property that their sum and difference are independent of each other, then the random variables have a joint Gaussian distribution (Kac [1939], Bernstein [1941], and see Kagan et al. [1973] for many much more general results). Since *R*_*M*_ is (half) the difference and 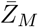 is (half) the sum of *Z*_*M*_ and *V*_*M*_, and *Z*_*M*_ and *V*_*M*_ are independent, if we assume that also *R*_*M*_ and 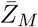 are independent, then both are therefore jointly Gaussian.

We have shown the following:

**Proposition 1**. *Suppose that* 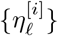 *are independent for all ℓ and i, that the distribution of* 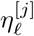 *only depends on ℓ, and that X*_*ℓ*_ *are independent of* 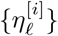. *If the Mendelian sampling term R*_*M*_ *is independent of the midparent value* 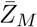, *then R*_*M*_ *and* 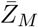 *are jointly Gaussian*.

Since models can be easily set up for which trait distributions are not Gaussian, the implication of this is that for such models, independence of the Mendelian sampling term is not likely a good assumption. Note that the proof does not depend on independence of the *X*_*ℓ*_: indeed, alleles may be inherited together, although in practice this would quickly create dependencies between nearby allelic effects. Also note that it is *not* usually true that the components – for instance, 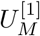 and 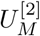 – are independent of each other.

As we discuss more later, the presence of a large effect polymorphic allele makes *R* and 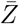 dependent: consider the extreme case where 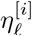 are i.i.d. with variance 1 for *ℓ* ≥ 2, while 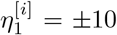 with probability 1/2 each, and 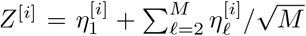. Then in the limit *R* depends strongly on parental trait values: if *Z*^[1]^ ≈ *Z*^[2]^ and so 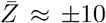 then most likely 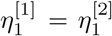 and so *R* is Normal with variance 1/2; but if 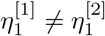 (and so 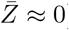) then *R* is close to +5 or −5 with equal probability.

## 4 Abnormal quantitative genetics?

As observed in Turelli [2017], there are at least three interpretations of what is called “the infinitesimal model”. The maximalist of these, what Turelli [2017] calls the Gaussian-population approximation, assumes a large number of small-effect loci, parental trait-independent Gaussian Mendelian sampling terms (as rigorously justified in Barton et al. [2017, 2022]), and, moreover, that population trait values are in deterministic mutation-selection balance, with a Gaussian empirical distribution (*i*.*e*., the *fraction* of the population with trait value in [*z, z* + *dz*) is approximately 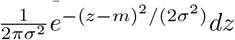 for suitable choices of the empirical mean and variance *m* and *σ*^2^). Leaving aside the question of whether or not some combination of effect sizes, dominance, and epistasis can justify non-Gaussian, trait-independent, Mendelian sampling terms, we might reasonably ask when (or if) deterministic mutation-selection balance is possible in a non-Gaussian model that retains independence of the Mendelian sampling terms. (Indeed, above we have put substantial restrictions on what models of inheritance might lead to such models.) We might also ask whether such a model might still reasonably approximate an additive model with non-Gaussian effects, and for the purpose of investigating this with simulations, some calculations will be helpful.

Concretely, consider a Moran process where an individual whose trait value is *Z* dies at rate *μ*(*Z*) and is replaced by the offspring of two randomly chosen individuals, i.e., a new individual with trait

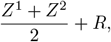

where *Z*^1^ and *Z*^2^ are independent draws from the empirical population distribution and *R* is an independent draw from the Mendelian sampling distribution (independent of *Z*^1^ and *Z*^2^, with distribution to be specified). Mutation-selection balance demands that that the distribution of individuals removed by death and the distribution of individuals added by birth are equal, *i*.*e*., that (*Z*^1^ + *Z*^2^)*/*2 + *R* is equal in distribution to a *μ*-weighted choice of *Z*. Taking the characteristic function of both these distributions, deterministic mutation-selection balance is therefore equivalent to

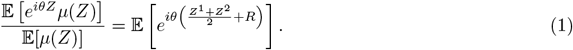

Now, we assumed that *Z*^1^, *Z*^2^, and *R* are independent, so we can write (1) as

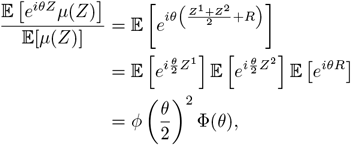

where 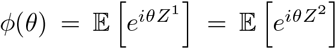 and Φ(θ) = 𝔼[*e*^*iθR*^] are the characteristic functions of *Z*^*j*^ and *R*, respectively. In what follows, we will explore some non-Gaussian solutions *ϕ*(*θ*) to (1).

### 4.1 The Neutral Case

First, consider the case without selection, so *μ* is a constant which we may take equal to 1. Then, our condition on the characteristic function simplifies to

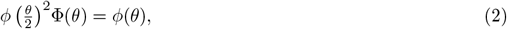

*i*.*e*., *ϕ*(*θ*) is a fixed point of *F*, where

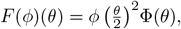

whereas any fixed point must be the limit of an arbitrary characteristic function, *φ*(*θ*), under repeated applications of *F, i*.*e*.,

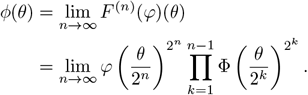

is a fixed point whenever this limit exists.

Since *φ* and Φ are characteristic functions, they are continuous and *ϕ*(0) = Φ(0) = 1, and thus both are non-zero in a neighbourhood of 0. In particular, *ψ*(*θ*) = Log *φ*(*θ*) and Ψ(*θ*) = Log Φ(*θ*) exist in some neighbourhood of 0 as well^1^. From the above, we have

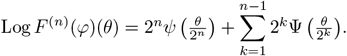

We briefly explore what possible limiting values are valid characteristic functions. We refer to the first term on the left hand side as the *reproduction term* and the remaining sum as the *noise term*, and only consider to the case when both have well-posed limits independent of the other.

#### Noise Terms

First, consider the noise terms. A necessary condition for 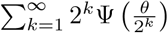 to converge is that 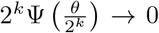; since Ψ(0) = 0, this implies that if it exists, Ψ^*′*^(0+) = 0. In particular, as we would expect, if *R* has a mean, it must be 0. If, on the other hand, Ψ(*θ*) is twice differentiable, so *R* has mean 0 and finite variance, say Ψ^*′′*^(0) = *η*^2^, then Taylor’s theorem tells us that for *θ* sufficiently small,

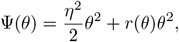

where *r*(*θ*) → 0 as *θ* → 0. Thus,

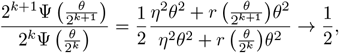

as *k* → ∞ and the sum converges by the ratio test. In this case,

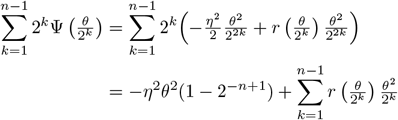

If we assume that *r*(*θ*) ≡ 0, so that the noise is *N* (0, *η*^2^), then the limiting distribution is *N* (0, 2*η*^2^). When *r*(*θ*) is non-zero, we don’t expect the higher order terms to vanish.

It is unclear what more can be said in general. If however, we specialize to the case when *R* is a stable law, we can explicitly compute the limiting noise term. We recall that if Ψ(*θ*) corresponds to a stable law (see *e*.*g*., Breiman [1992]), *then*

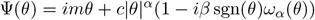

for *α* ∈ (0, 2], *β* ∈ [−1, 1], *c* ≥ 0, and *m* ∈ ℝ and

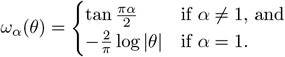

By our general remarks above, we can assume *m* = 0 and exclude the case *α* = 1, *β* = 0, for which Ψ^*′*^(0+) = *c*. We then have

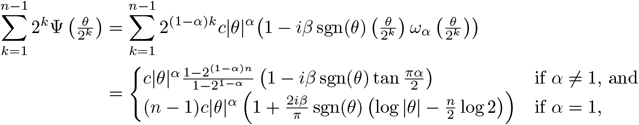

which converges as *n* → ∞ if and only if *α >* 1, in which case the limit is

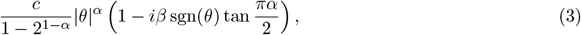

which corresponds to a 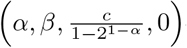 stable law.

#### Reproductive Terms

We now turn our attention to the limit

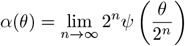

If it exists, then for all integers *k*,

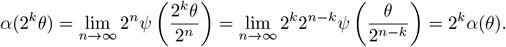

and, since *α*(0) = 0,

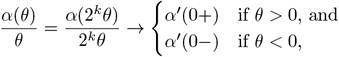

as *k* → −∞ if the limits exist (*i*.*e*., there are unique left-and right-hand limits at zero; this is the case if *e*.*g*., we require *α*(*θ*), and thus *ϕ*(*θ*), to be convex functions). We henceforth assume this.

Since *φ*(*θ*) is a characteristic function, it has the Hermitian property: 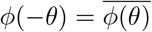, and thus 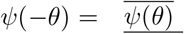 and 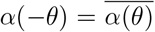. In particular, if one (and by extension both) limits exist, we must have 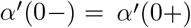. In this case, setting −*γ* + *im* = *α*^*′*^(0+), we have

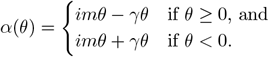

If *γ >* 0, then we recognize 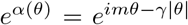 as the characteristic function of a Cauchy distribution with median 0 and median absolute deviation *γ*, whereas if *γ* = 0, then 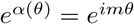 is the Fourier transform of a Dirac mass at *m, i*.*e*., the characteristic function for a monomorphic population with all individuals having trait value equal to *m*. (And, *γ <* 0 does not give a valid characteristic function.)

#### Blending inheritance redux

Note that the characteristic functions constructed above are solutions to (2) when transmission is noiseless, and the offspring have exactly the parental mean trait – *i*.*e*., blending inheritance. The solution 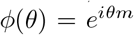 is exactly the reversion to the mean that Darwin’s detractors used to argue against the theory of evolution [Jenkin, 1867, Provine, 1971]. The solution 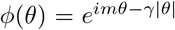 offers an amusing counter-factual: if a Lamarkian population could somehow achieve a Cauchy-distributed trait distribution, it would maintain that diversity indefinitely.

More generally, one will get convergence if the noise is a mixture of components with finite variance and stable laws as above, whereas the population distribution will be a mixture of a (possibly degenerate) Cauchy distribution and such noise terms. Note that in the presence of noise, a pure Cauchy-distributed solution is *not* possible in the neutral case.

### 4.2 Stabilizing Selection

Unfortunately, once selection is included, (1) is less amenable to the sort of simple arguments above. We will content ourselves with presenting two examples of stabilizing selection, one already familiar to quantitative genetics, the other a Cauchy solution obtained by trial-and-error, which illustrates how selection can stabilize an otherwise unviable solution.

#### Gaussian Solution

First, consider the case when 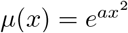. In this case there is a Gaussian solution: suppose *Z* is Gaussian with mean *m* and variance *σ*^2^, so 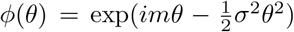. Moreover, provided 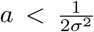, we can explicitly compute 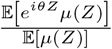: if *R* is Gaussian with mean zero and variance *η*^2^, then equation (1) becomes

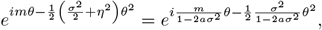

which, equating real and complex components, has a solution provided *m* = 0 and

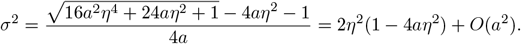

Note that we do not require very weak selection to guarantee a Gaussian steady state, only that 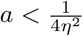. However, this does not imply that the transient distribution is also Gaussian, as is sometimes assumed [e.g., Lande, 1976].

##### 4.2.1 Cauchy Solution

Almost uniquely among the stable laws, the Cauchy distribution has a probability density function expressible in elementary functions: for a Cauchy random variable with median (location) *x*_0_ and mean absolute deviation (scale) *γ*, the density is

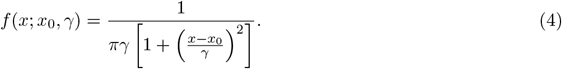

Armed with this density function, we can explicitly compute the right hand size of (1), assuming that

i. the population is in mutation-selection balance corresponding to a Cauchy density with median 0 and scale *γ >* 0,
ii. selection acts as 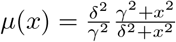 for some *δ > γ >* 0, and
iii. the Mendelian sampling term *R* is given by a Cauchy(0, *δ* − *γ*) random variable (*i*.*e*., corresponding to a stable (1, 0, *δ* − *γ*, 0) law).

Under these assumptions, (1) is

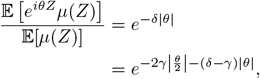

which we recognize as being equal to 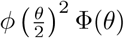, where 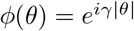 and 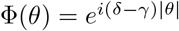 are the characteristic functions of the population density and the Mendelian sampling term, respectively, giving a.non-Gaussian solution to (1) with both noise and selection.

As remarked above, we found this solution by inspection and trial-and-error; it is interesting to ask if any similar examples can be constructed for other stable laws. The lack of an elementary probability density function unfortunately rules out the naïve approach used here.

## 5 The difficulty with an abnormal quantitative genetics

Above we have seen that (a) in principle, a mutation-selection balance is possible with a Cauchy empirical distribution and Cauchy Mendelian noise, but (b) such a model cannot derive from a simple model of additive loci. To that end, it’s instructive to see where the argument of Barton et al. [2017] breaks down if instead of assuming bounded allele effects of order 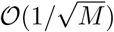, we assume that the allele at locus *ℓ* in individual *j* contributes 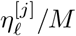, where the 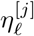 are independent Cauchy(0, *γ*_*ℓ*_) random variables. In this case, the typical effect size at locus *ℓ* is of order 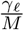, which is asymptotically smaller than the allele effects assumed in Barton et al. [2017], but large effect alleles are more likely.

The crucial step in the derivation of the infinitesimal model in Barton et al. [2017] is to show that, following the arguments in Fisher [1918], knowing an individual’s trait value gives vanishingly little information about the contribution of any given locus as *M*→ ∞, which in turn allows one to show that the within-family Mendelian segregation variance is independent of the parental traits. Here, we will consider how conditioning an individual’s trait value affects the law of a single allelic contribution in the Cauchy setting described above. A similar argument can be made for any *α*-stable distribution, but the lack of explicit probability density functions for generic *α* ∈ (0, 2) adds complexity to the argument without adding any insight.

Consider the simplest possible case of an ancestral population, of *N* individuals, where the *j*^th^ individual has trait value

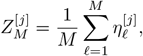

where the 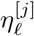 are independent across individuals and across loci. By the additive property of Cauchy random variables, 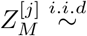, where

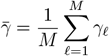

is the mean of the scale parameters.

To see what knowing the parental trait value 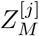 tells us about the contribution of any given locus, 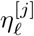, we follow the arguments in Fisher [1918] and Barton et al. [2017] using Bayes’ Theorem. Proceeding informally, with the probabilities indicating probability density functions, we have

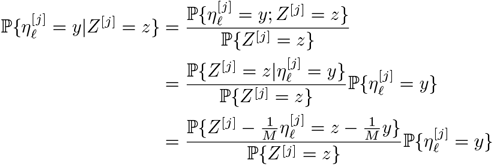

Now, 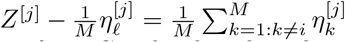 is Cauchy-distributed with scale 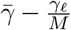. Using the probability density function for a Cauchy distributed random variable (4), a straightforward computation gives

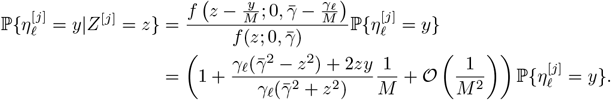

Noting that 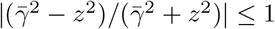 and 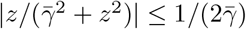 (since it is maximized at 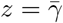), so

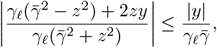

we have that

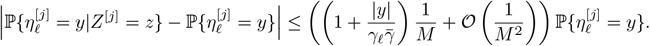

In the case of bounded allelic effects, the error introduced by approximating the effect-size distribution conditional on the trait value by the unconditioned distribution is proportional to |*Z*^[*j*]^|, so that extreme trait values are most informative of the size of allelic effects, whereas here, the bound is independent of the trait value, and the error is maximized when the trait value is approximately equal to the mean absolute deviation of its distribution, *i*.*e*., the median value of |*Z*^[*j*]^|.

The argument above shows for a typical locus, the effect of conditioning on the trait gives negligible information about the contribution from that site. Recalling that

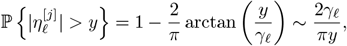

we see that for any *ϵ >* 0, and large values of *M*,

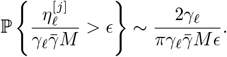

so the probability the error is larger than *ϵ* at any given locus decays rapidly as *M*→ ∞.

If, however, we consider not any one locus, but rather all loci, we get a different picture. The variables 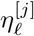 are independent, and thus so are the Bernoulli random variables

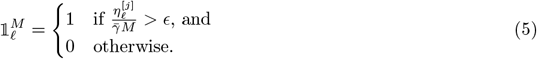

Now, since

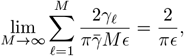

the number of loci at which the magnitude of the effect size exceeds *Mϵ*, which is 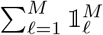, converges weakly as *M* → ∞ to a Poisson random variable with mean 2*/*(*πϵ*) (see *e*.*g*., Theorem 2.6.1 in Durrett [2005]). In particular, for large values of *M* and fixed *ϵ >* 0,

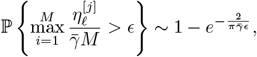

which approaches 1 as *ϵ* → 0, while the expected number of *M* alleles such that 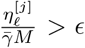 is asymptotically 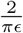.

In particular, there is at least one locus at which knowing the parental trait gives one information about the effect size (and there are on the order of 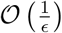 that give “an *ϵ* of information”), and these large effect loci in turn give rise to correlations between parental traits and the deviation of the offsprings’ traits from the midparent (i.e., the Mendelian sampling terms). However, the first calculations above suggest that for “most” loci, this is not true, and so perhaps a reasonable picture of Mendelian sampling terms in an additive model with Cauchy-distributed effects might be “some large Mendelian alleles plus Gaussian noise”. We now turn to simulation, to investigate.

## 6 Simulations

In simulation we can easily include realistic recombination (instead of unlinked loci) and selection. We used SLiM v4 [Haller and Messer, 2022] to simulate a discrete-time approximation to the continuous-time Moran model described above as follows. Individuals are diploid, hermaphroditic, and sexual; the genome is of length 10^8^bp with a uniform recombination rate of 10^−8^ crossovers per bp; mutations occur at a rate of 10^−9^ per bp. Each new mutation is independently assigned an “effect” drawn from the “effect size distribution” (and will be either Gaussian or Cauchy). The trait of each individual is determined additively by summing the effects of all mutations they carry across both chromosomes. Let *μ*(*z*) be the death rate of an individual with trait *z*, and let *dt* be a small constant (we took *dt* = 0.01). In all plots, one time unit is approximately one generation. Then, at each time step each individual dies with probability 1 − exp(−*dtμ*(*z*)); if there are *k* such individuals chosen to die in a given time step, then we choose *k* individuals uniformly and without replacement to produce new offspring; for each such reproduction event the parent chooses a mate uniformly at random from the population. This maintains the population at a fixed size, *N*. (Note that individuals may self and individuals chosen to die may also reproduce, but in a large population these details should be unimportant.)

We first consider the neutral case (i.e., with *μ* ≡ 1). As shown in Figure 3, a simulation with a Cauchy effect size distribution, unsurprisingly, has a distribution of trait values with more extreme values and a rougher path of population median trait value over time than does a simulation with a Gaussian effect size distribution. Indeed, the sum of all effects along the path from the start of the simulation to an arbitrary individual will approximate either Brownian motion (in the Gaussian case) or a Cauchy process (in the Cauchy case). Figure 4 shows how the Mendelian sampling term depends on midparent value. In the Gaussian case, the Mendelian noise is Gaussian and independent of midparent value, as shown by the horizontal lines at 1.0 in Figure 4C. On the other hand, in the Cauchy case we can see from either Figure 4D or F that Mendelian noise depends strongly on midparent trait, as we would expect in the presence of several large-effect alleles.

**Figure 3.**
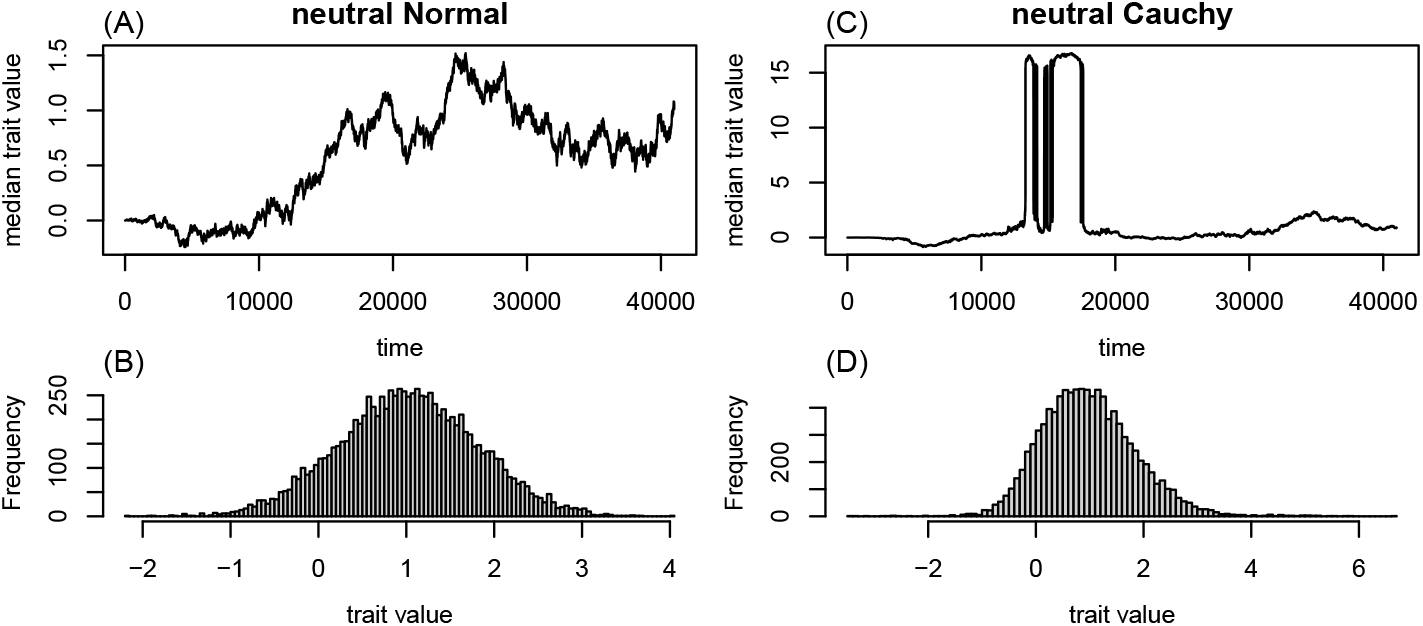
Median trait values over time **(A and C)** and distributions of final trait values **(B and D)** in neutral simulations with Gaussian **(A and B)** and Cauchy **(C and D)** effect size distributions. Simulations were begun with no genetic variation, had populations of size *N* = 1000, a genome of length *L* = 10^8^ bp and mutation rate of *μ* = 10^−9^ per bp per generation; the effect size distributions were Normal with mean 0 and standard deviation 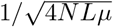 and Cauchy with center 0 and scale 1*/*(4*NLμ*), respectively.

**Figure 4.**
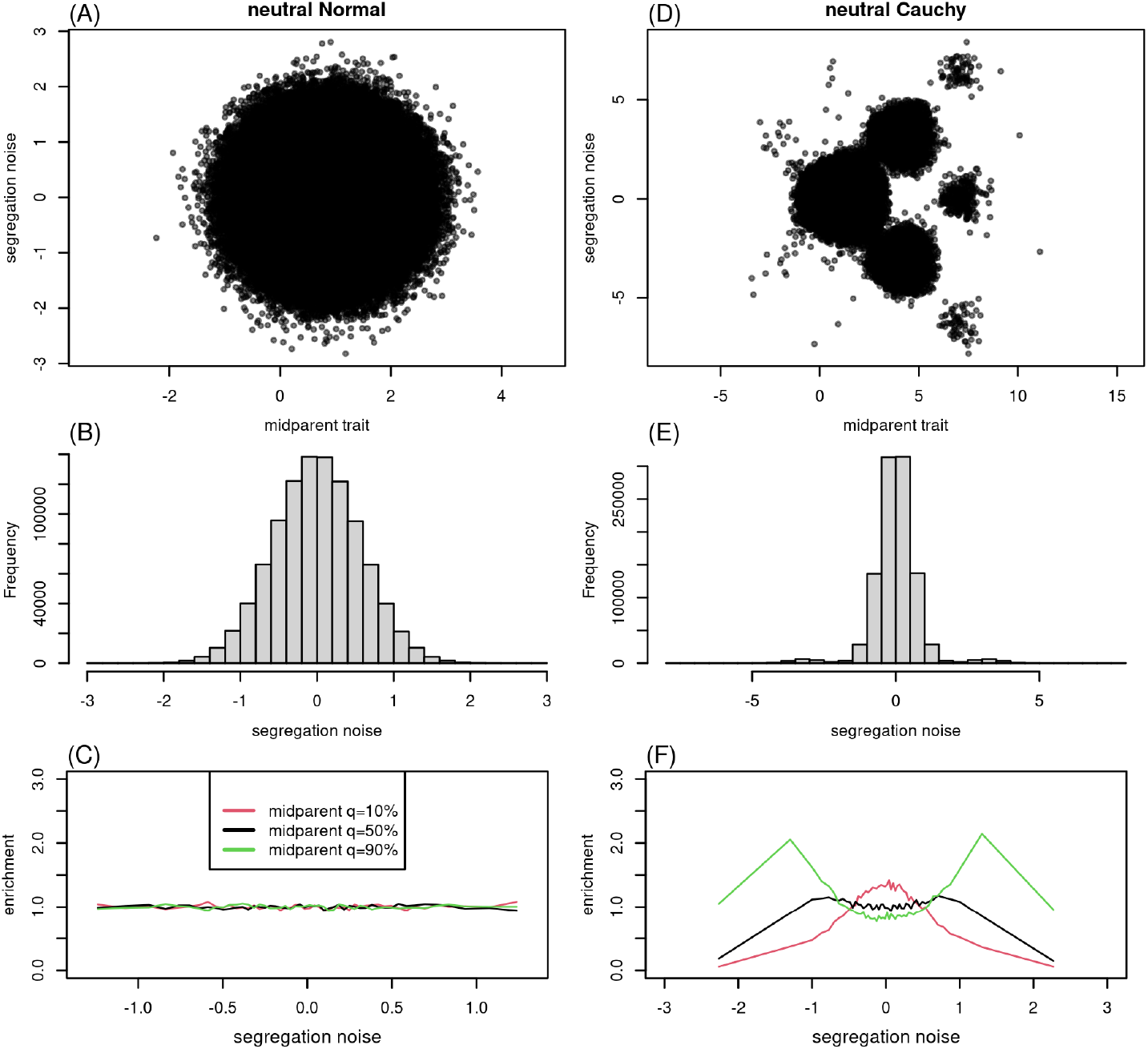
The distribution of Mendelian sampling terms with Gaussian **(A-C)** and Cauchy **(D-F)** effect size distributions, from the same simulations shown in Figure 3, for all individuals born during the last 1,000 time units of the simulation. **(A**,**D)** Mendelian noise (offspring trait minus midparent) plotted against midparent. **(B**,**E)** Histograms of the Mendelian sampling term. **(B**,**E)** Enrichment in quantiles of the Mendelian sampling term conditional on midparent value. To compute the latter, let *q*_0_, *q*_2_, *q*_4_, …, *q*_98_, *q*_100_ divide the distribution of the Mendelian sampling term into 50 bins with roughly equal numbers in each (i.e., the quantiles), let *p*_*k*_(*a, b*) denote the proportion of the offspring of parents with midparent value is in [*a, b*) whose Mendelian sampling term is in [*q*_*k*−1_, *q*_*k*_), and let *p*_*k*_ be the proportion of all offspring with Mendelian noise in [*q*_*k*−1_, *q*_*k*_) (so, *p*_*k*_ ≈ 0.02). Then, each line shows *p*_*k*_(*a, b*)*/p*_*k*_ plotted against the midpoint of the corresponding quantile bin (*q*_*k*−1_, *q*_*k*_), for (*a, b*) chosen to span the 5% of midparents centered on: (red) the 10% quantile, (black) the median, and (green) the 90% quantile of midparent value.

We next compared two situations of stabilizing selection, motivated by the results in Section 4.2: (1) the effect size distribution was Gaussian with mean zero and standard deviation 1.0 and 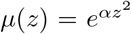; and (2) the effect size distribution was Cauchy, centered at zero and with median absolute deviation *δ*− *γ* and *μ*(*z*) = (*δ/γ*)(*γ* + *z*^2^)*/*(*δ* + *z*^2^) with *δ* = 0.5 and *γ* = 0.2. Figures 5 and 6 show that the picture is largely unchanged by stabilizing selection except that, unsurprisingly, the median trait no longer wanders unconstrained (but still shows substantially larger fluctuations under the Cauchy model).

**Figure 5.**
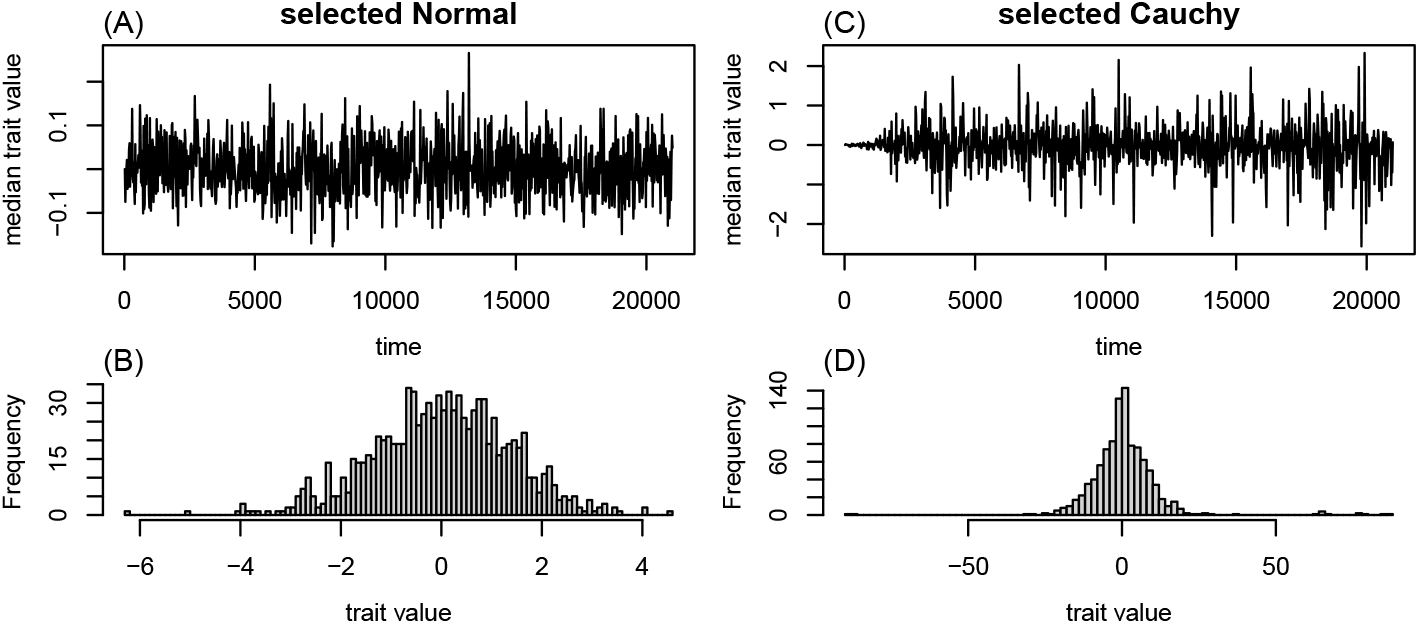
As in Figure 3, except under a model of stabilizing selection (see text for details).

**Figure 6.**
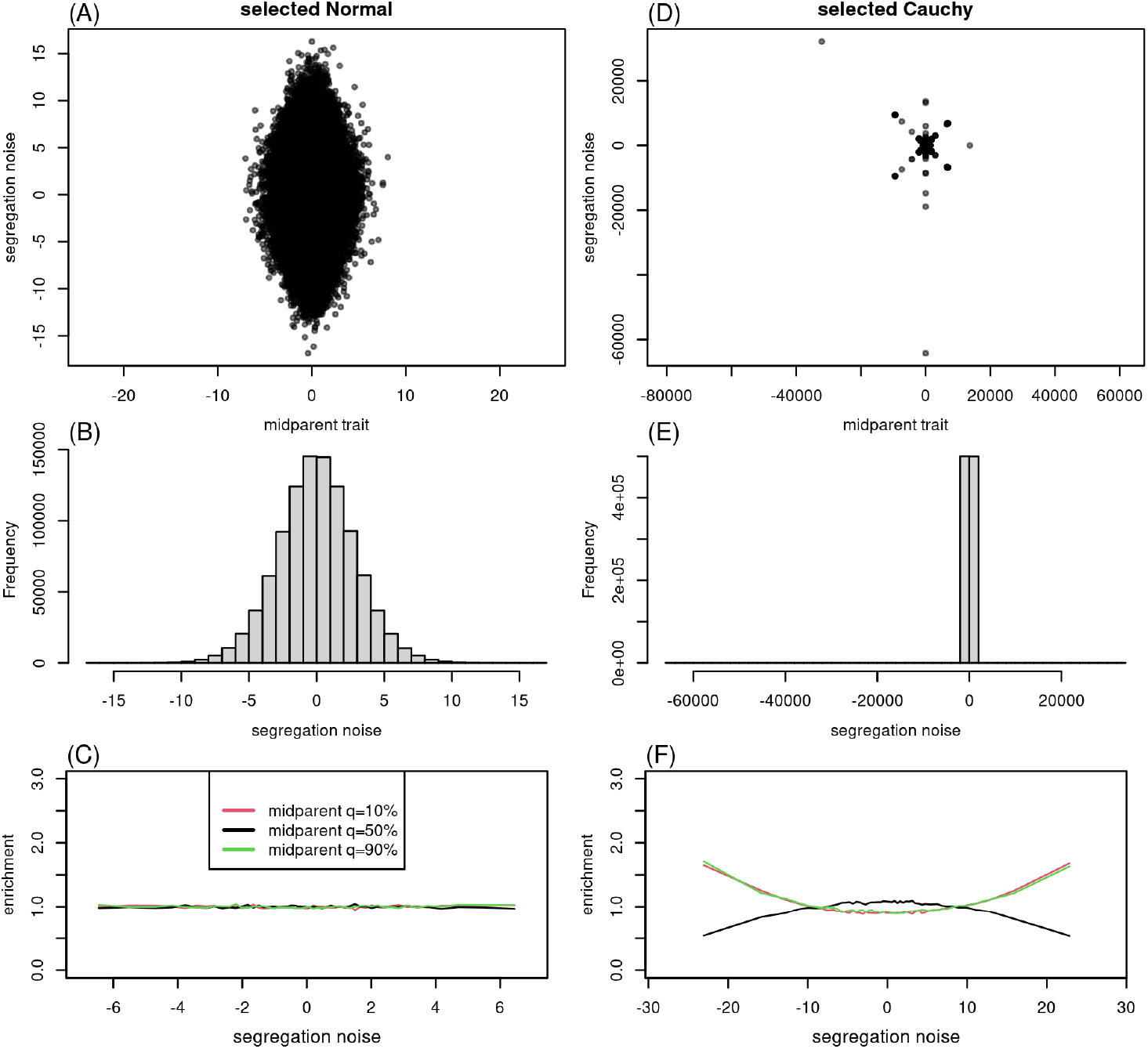
As in Figure 4, except under a model of stabilizing selection (see text for details).

However, the two models look much more similar when considering only alleles of small effect. Figure 7 shows that midparent values and Mendelian sampling terms computed using only mutations with effect size less than 0.2 (excluding less than 1% of mutations) are quite close to independent. The result does not depend strongly on the cutoff chosen, and are similar even if less 0.1% of mutations are excluded.

**Figure 7.**
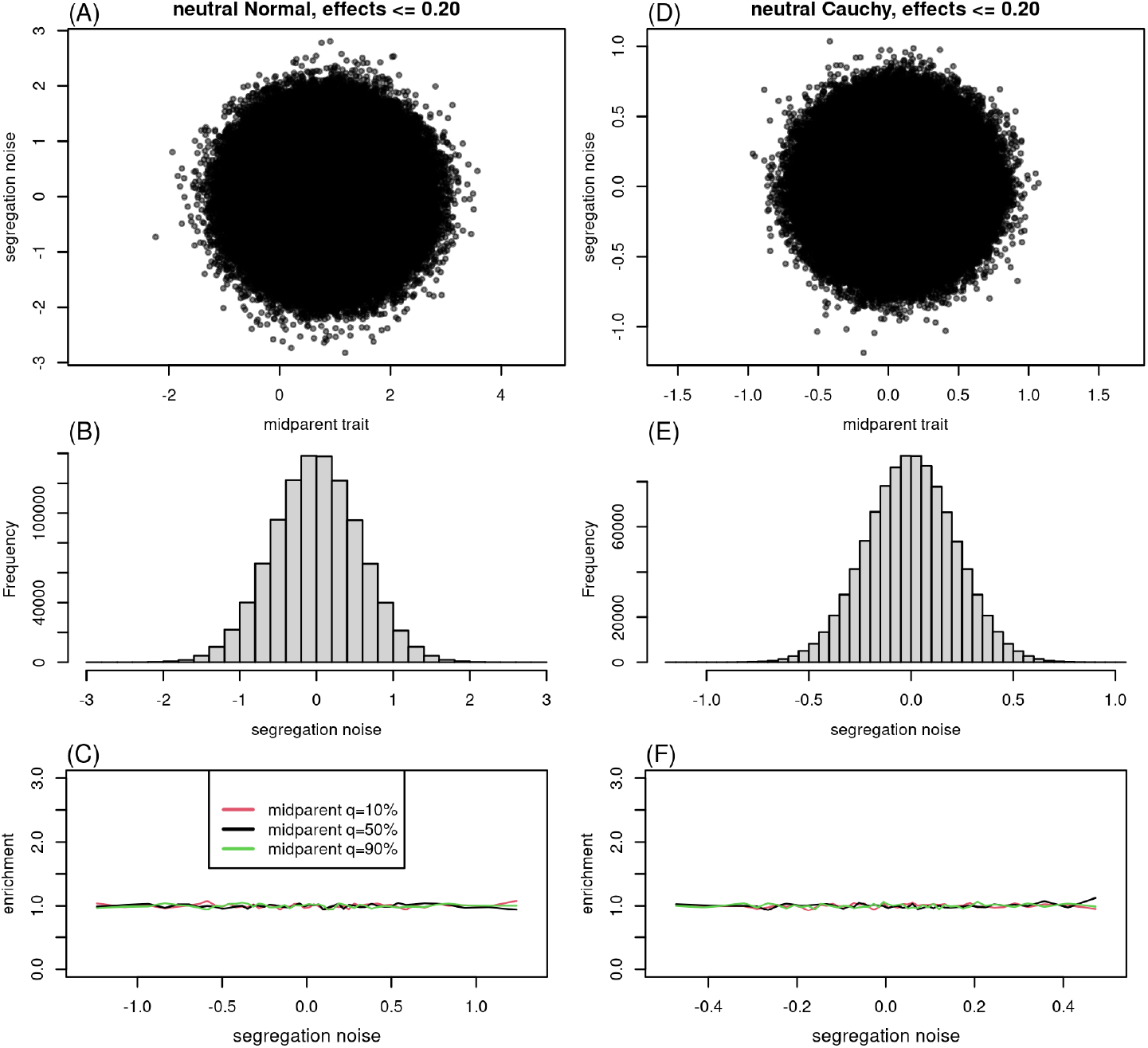
As in Figure 4, except computed using only mutations with absolute value effect size less than 0.2. The same quantities for the simulations of Figure 6 with stabilizing selection are shown in Figure S2.

## 7 Discussion

A substantial challenge for the field is to develop tractable models of trait evolution that include contributions of both “large” and “small” effect loci. In practice, breeders have long embellished the “animal model” (essentially, the infinitesimal model) to predict traits by combining both effects of known large effect loci and cumulative effects of many unknown loci as mediated by a relatedness matrix [e.g., Fernando and Grossman, 1989, Teissier et al., 2018, Bernardo, 2014, Rice and Lipka, 2019]. We can study any model with simulation, but the way forwards to theoretical understanding seems murky.

Further analysis of the infinitesimal model with other noise distributions, as we have begun here, could be interesting, e.g., to see how changing the distribution of the Mendelian sampling term affects the rate of adaptive evolution, levels of genetic load, the rate of fixation, and the strength of linked selection. However, we have seen that the assumption of independence in the infinitesimal model implies a sort of “blending inheritance” for the large effect alleles, and so such work must be accompanied by simulations to see if the predictions are borne out under concrete models of genetic inheritance. Another open question is whether there is some set of realistic assumptions (perhaps involving epigenetic effects, pleiotropy, and/or environmental interactions with genetic effects) which leads to these non-Gaussian infinitesimal models. Finally, it may well be that there is a clever way to set up a trait-only model of evolution that is both insensitive to the underlying genetics and does not rely on the Gaussian central limit theorem.

Finally, we note that above we’ve most often used the Cauchy distribution not because it seems most realistic (indeed data suggests the Lévy stable distribution with *α* = 3*/*2 might be better) but for mathematical convenience.

## Acknowledgements

Thanks go to Nate Pope for useful discussion and to Gregor Gorjanc for insight into modern uses of the animal model and genomic selection. This project was started on a visit by TLP to the University of Oregon that was partially supported by the CNRS PEPS grant “Jeunes Chercheuses et Jeunes Chercheurs”.

---

**Figure S1:**
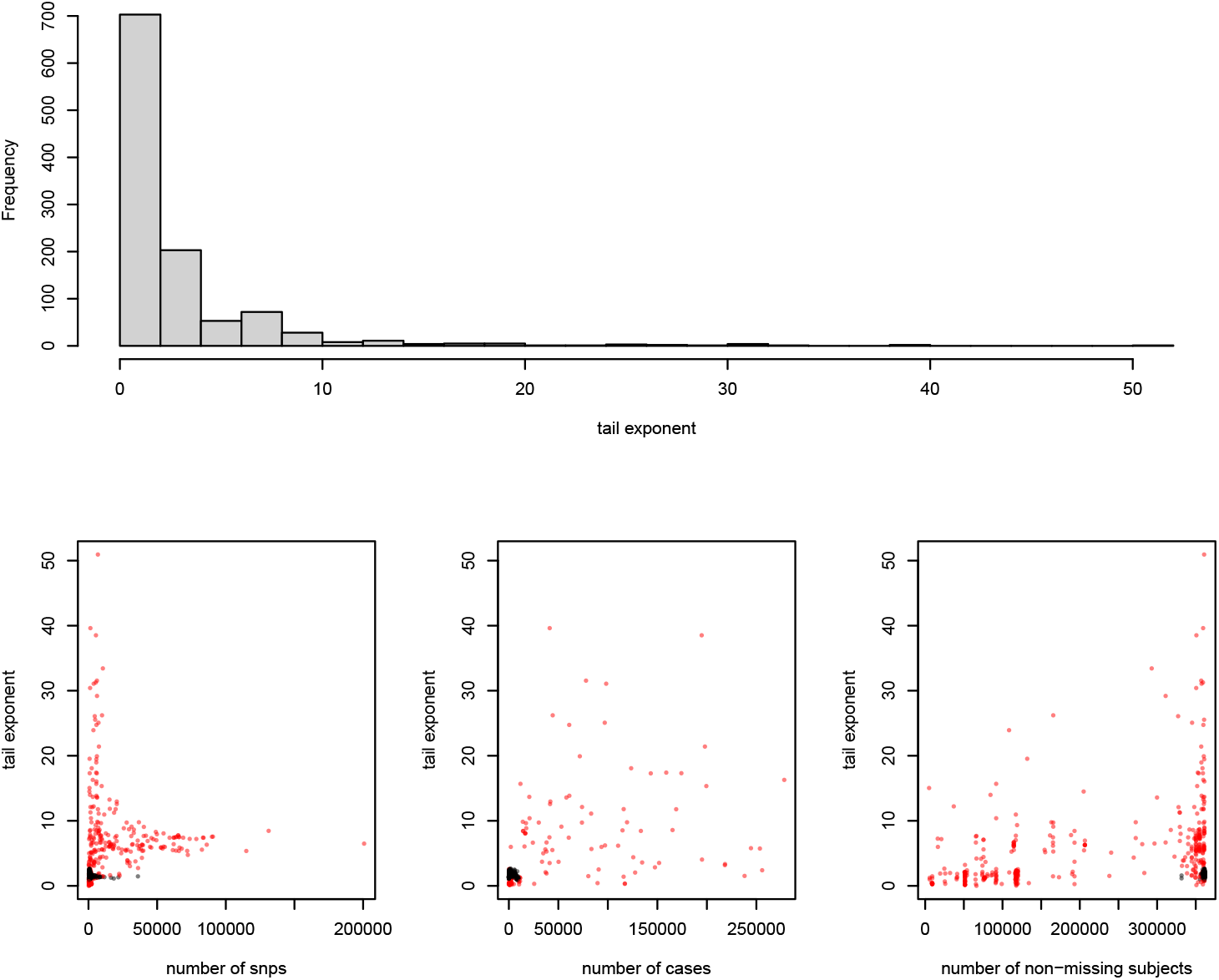
Estimated values of the tail exponent, *α*, for 1108 illness-related binary phenotypes, including the 333 phenotypes removed in filtering, which are shown as red points in lower plots. **(A)** distribution of values; and plotted against **(B)** number of SNPs, **(C)** number of cases, and **(C)** number of non-missing subjects.

**Figure S2:**
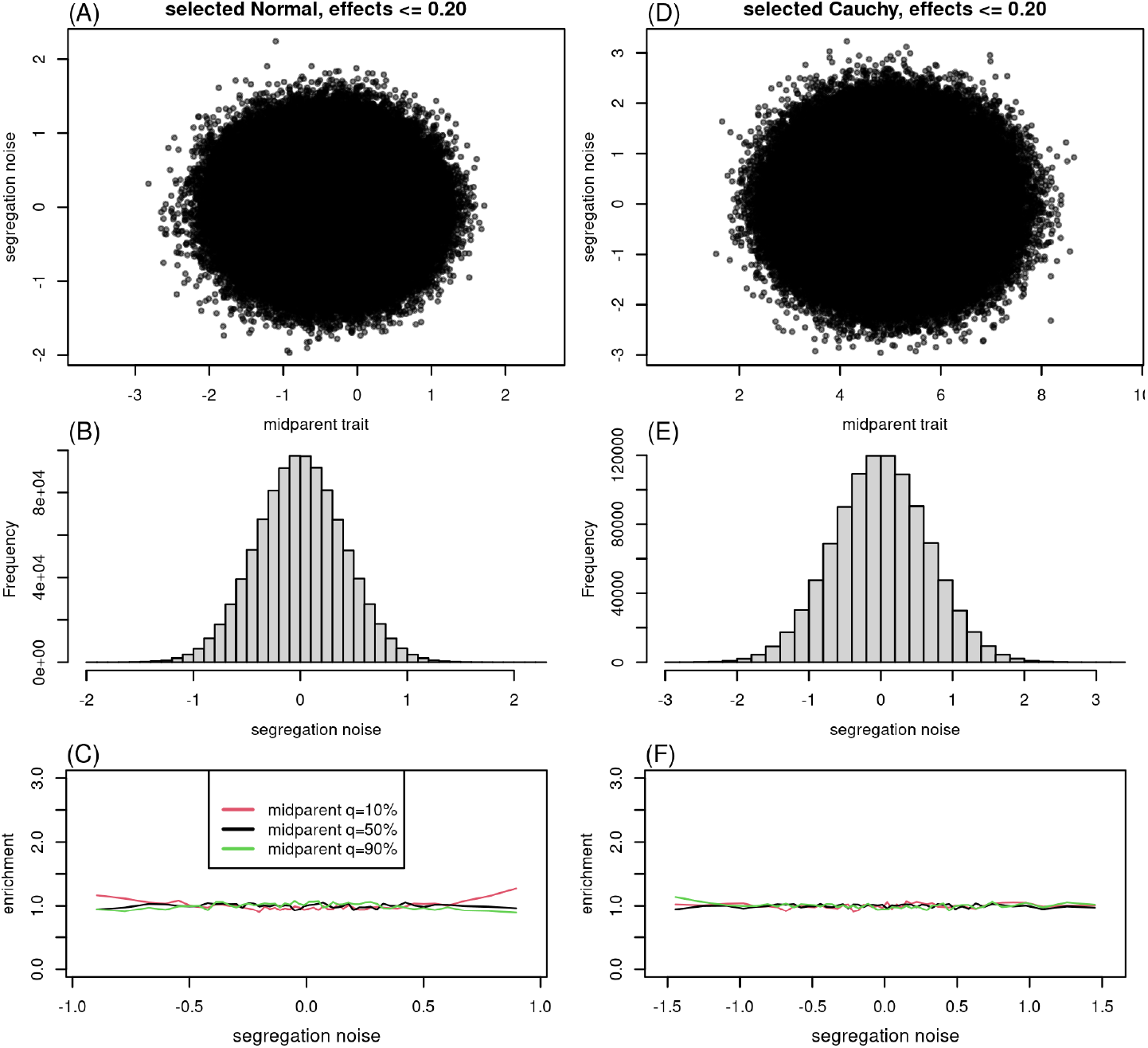
As in Figure 4, except computed using only mutations with absolute value effect size less than 0.2.

## Footnotes

1 We use Log to denote the principal value of the complex logarithm, for which the imaginary part always lies in the interval (−*π, π*].

## Notes

### Competing Interest Statement

The authors have declared no competing interest.

